# Chronic behavioral manipulation via orally delivered chemogenetic actuator in macaques

**DOI:** 10.1101/2021.08.03.454990

**Authors:** Kei Oyama, Yukiko Hori, Yuji Nagai, Naohisa Miyakawa, Koki Mimura, Toshiyuki Hirabayashi, Ken-ichi Inoue, Masahiko Takada, Makoto Higuchi, Takafumi Minamimoto

**Affiliations:** Department of Functional Brain Imaging, National Institutes for Quantum and Radiological Science and Technology, Chiba 263-8555 Japan; Systems Neuroscience Section, Primate Research Institute, Kyoto University, Inuyama, Aichi 484-8506, Japan; PRESTO, Japan Science and Technology Agency, Kawaguchi, Saitama, Japan

## Abstract

The chemogenetic technology referred to as designer receptors exclusively activated by designer drugs (DREADDs) offers reversible means to control neuronal activity for investigating its functional correlation with behavioral action. Deschloroclozapine (DCZ), a recently-developed highly potent and selective DREADDs actuator, displays a capacity to expand the utility of DREADDs for chronic manipulation without side-effects in nonhuman primates, which has not yet been validated. Here we investigated the pharmacokinetics and behavioral effects of orally administered DCZ in macaque monkeys. Pharmacokinetic analysis and positron emission tomography (PET) occupancy examination demonstrated that oral administration of DCZ yielded slower and prolonged kinetics, and that its bioavailability was 10-20% of that in the case of systemic injection. Oral DCZ (300-1000 μg/kg) induced significant working memory impairments for at least 4 h in monkeys with hM4Di expressed in the prefrontal cortex. Repeated daily oral doses of DCZ consistently caused similar impairments over two weeks without discernible desensitization. Our results indicate that orally delivered DCZ affords a less invasive strategy for chronic but reversible chemogenetic manipulation of neuronal activity in nonhuman primates, and this has potential for clinical application.

## Introduction

The chemogenetic tool referred to as DREADDs (designer receptors exclusively activated by designer drugs) has been widely used for controlling neuronal activity and animal behavior in many species, including rodents and nonhuman primates [1]. By combining mutated muscarinic receptors (excitatory hM3Dq and inhibitory hM4Di) with specific actuators such as CNO (clozapine-N-oxide), these chemogenetic techniques offer a means to remotely manipulate the activity of a specific neuronal population for minutes to hours [2]. In addition to the acute control, repeated administration of the actuators enables chronic chemogenetic manipulation of neuronal activity for days to weeks [3–10], and thus has potential therapeutic applications. For example, chronic and reversible inhibition of neuronal activity can be an effective means for seizure control and epilepsy treatment [11]. For this purpose, oral delivery of agonists seems to be suitable, as it typically produces relatively long-lasting effects compared to systemic (i.e., intramuscular and intravenous) injections [12]. Besides, oral administration is non-invasive and relieves the subject from stress during chronic treatment. It is also beneficial for neuroscience research aimed at long-term control of neuronal activity in freely-moving animals without restraint or stress. However, recent studies have reported that CNO has low permeability into the brain, and that its metabolite, clozapine, activates not only DREADDs but also endogenous receptors and transporters, leading to potential off-target effects [13,14]. To circumvent this issue, alternative selective DREADDs agonists have been explored and validated [15–17].

Deschloroclozapine (DCZ) is one of the 3rd-generation agonists for muscarinic DREADDs that represents high permeability into the brain and is highly potent and selective to DREADDs, as compared to prior agonists such as CNO and C21 [17]. Importantly, low doses of DCZ exert chemogenetic effects with rapid onset that last several hours in both rodents and nonhuman primates [17]. This indicates that DCZ is a promising chemogenetic actuator that can be used for a wide variety of objectives. In particular, its superior efficacy and brain permeability permit its use for oral administration. In fact, it has been demonstrated that orally delivered DCZ effectively induces behavioral changes in marmosets [18]. However, the long-term chemogenetic action of DCZ remains unclear, specifically in terms of the duration of a single-dosage effect and the impact of repetitive doses over several days to weeks. These will provide knowledge sufficient for designing chronic chemogenetic experiments and, in addition, for developing potential clinical therapeutic applications.

Here we investigated the pharmacokinetics and in vivo occupancy of orally administered DCZ to determine a dose range suitable for DREADDs studies in monkeys. We then demonstrated that oral DCZ administration induced a chemogenetic behavioral effect lasting at least more than 4 h in macaque monkeys with hM4Di expressed in the prefrontal cortex. We also found that repetitive daily administrations of DCZ yielded consistent behavioral effects over weeks without apparent sign of desensitization.

## Results

### Pharmacokinetics of orally administered DCZ

We first conducted pharmacokinetic studies in three naïve monkeys to estimate the time course of the chemogenetic effects of DCZ via oral administration. In our previous study, intramuscular administration of DCZ at 100 μg/kg provided a sufficient level of DCZ concentration in CSF to activate hM4Di receptors for 2 h, during which it gave rise to behavioral deficits in monkeys via hM4Di activation [17]. Given that oral administration of clozapine, an analogue of DCZ, results in relatively lower bioavailability and slower onset of action compared to intravenous administration [23], we first examined DCZ concentrations in the plasma and CSF for 4 h following oral DCZ administration (300 μg/kg), as well as intramuscularly (100 μg/kg) as reference. Oral DCZ administration yielded a maximum DCZ concentration in plasma at 60 min after administration, followed by a gradual decrease (Fig. 1a, orange). By contrast, intramuscular administration increased plasma DCZ concentration at 30 min, which then decreased monotonically (Fig. 1a, green). Both oral and intramuscular administrations produced relatively stable DCZ concentrations in CSF for at least 4 h (>2 nM and >5 nM; Fig. 1b, green and orange, respectively), which were sufficient concentrations to activate hM4Di DREADD (EC50 of 0.081 nM) [17]. DCZ was not detected in plasma or CSF 24 h after administration. We also examined the concentration of possible metabolites of DCZ, C21 and DCZ-N-oxide in plasma; however, metabolites were not detected, confirming the metabolic stability of DCZ (Fig. 1a) [17]. Taken together, these results suggest that the oral administration of DCZ produces a relatively long-lasting pharmacokinetic profile without significant metabolite production and a lower bioavailability, about 10–20% of that of intramuscular administration.

**Figure 1.**
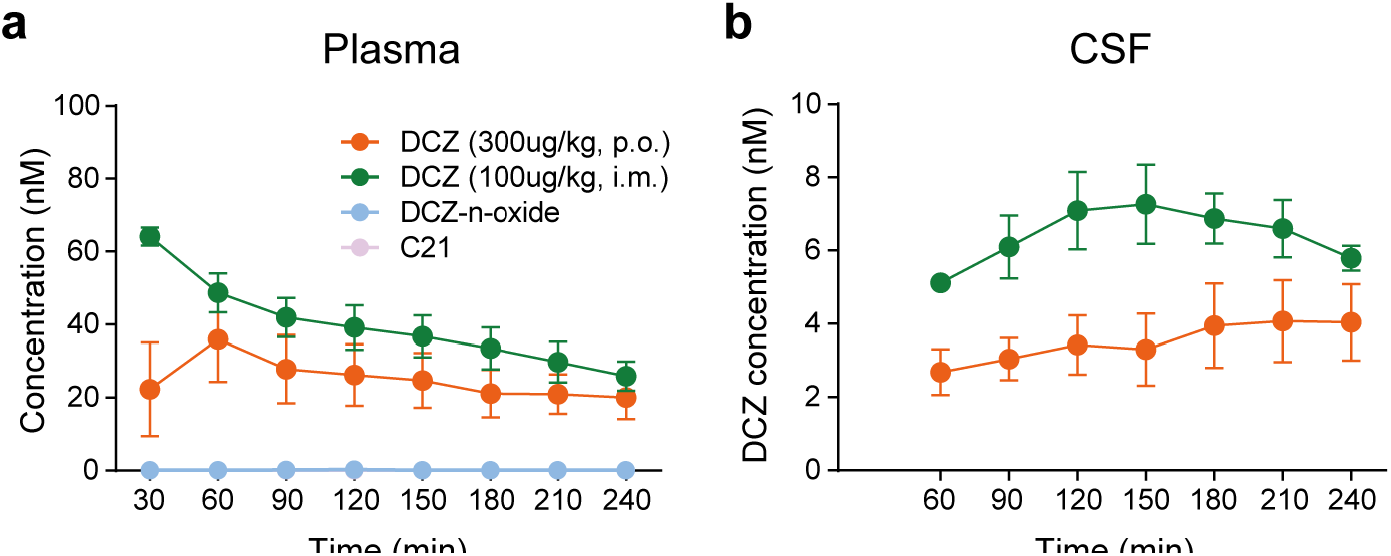
Time concentration profiles of DCZ by oral and intramuscular administration. A, Time-course of DCZ concentration in the plasma (**a**) and CSF (**b**) following intramuscular injection of 100 μg/kg (green) and oral administration of 300 μg/kg (orange) DCZ. Concentration of major metabolites of DCZ (DCZ-n-oxide and C21, light blue and purple, respectively) following oral administration of 300 μg/kg DCZ are also shown (data for C21 were behind those of DCZ-n-oxide as neither were detected). Data were collected from 3 monkeys (#226, #248, and #249).

### Measuring in vivo occupancy of hM4Di-DREADD by oral DCZ administration

Next, to determine an effective and safe dose range for oral DCZ administration, we measured hM4Di receptor occupancy by DCZ — the relationship between DCZ dose and the degree of hM4Di receptor occupation (Fig. 2a). This was important because a higher agonist dose will afford a stronger chemogenetic effect, but may risk higher off-target effects [24]. In our previous study, a series of PET studies were used to determine the optimal DCZ dose that would yield a 50% to 80% occupancy of hM4Di, which was 30 to 100 μg/kg for systemic administration (Fig.2c, see [17] for details). Given that the bioavailability of oral administration would be 10-20% of that of systemic administration (Fig. 1), we estimated that oral DCZ doses achieving 50% and 80% hM4Di occupancy would be around 300 and 1,000 μg/kg, respectively. We performed a PET occupancy study using a monkey that had received injection of a viral vector encoding an hM4Di gene into the unilateral amygdala. Baseline PET imaging with radiolabeled DCZ ([^11^C]-DCZ) confirmed that the hM4Di expression at the injected side of amygdala was indicated by an increased radioligand binding (Fig 2b, baseline). Subsequent [^11^C]-DCZ PET scans were performed after oral pretreatments (i.e., oral administration of vehicle, 300, or 1,000 μg/kg of non-radiolabeled DCZ). Consistent with our previous findings, specific binding of [^11^C]DCZ at the hM4Di-expressing amygdala region was diminished with increasing doses of DCZ (Fig. 2b). Occupancy was determined as a reduction of specific radioligand binding (BPND) at the target region over the control side relative to baseline (Fig. 2b, see Materials & Methods for details). As we predicted, the hM4Di occupancies with 300 and 1,000 μg/kg oral administration were about 40% and 90%, respectively (Fig. 2c). The dose for 50% occupancy (EC_50_) was 285 µg/kg, which corresponded to about a 10-fold dose of systemic administration and was roughly consistent with the bioavailable profile shown in our pharmacokinetic study described above. Based on these results in pharmacokinetic and occupancy studies, we determined the effective and safety dose range for oral DCZ administration to be 300–1,000 μg/kg.

**Figure 2.**
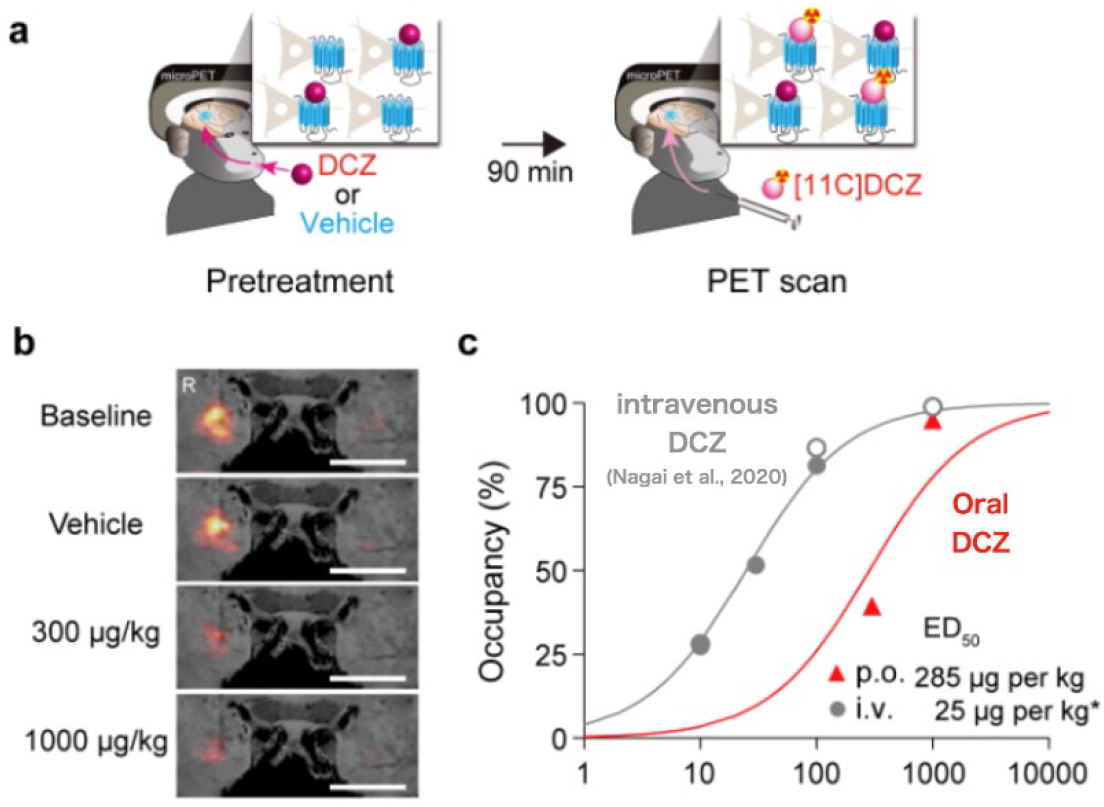
PET visualizes the occupancy of orally administered DCZ for hM4Di. **a**, Schematic illustrations of occupancy study by PET. Monkeys underwent [^11^C]DCZ-PET scan 90 min after p.o. administration of nonradiolabeled DCZ or vehicle. **b**, Coronal sections of parametric [^11^C]DCZ-PET image of specific binding (BPND) overlaying MR image of a monkey expressing hM4Di in the putamen. Scale bars represent 5 mm. **c**, Occupancy of hM4Di plotted as a function of DCZ. Red triangles represent occupancy by oral DCZ doses obtained from monkey # 237. Curves are the best-fit Hill equation for the data. ED50 indicates the agonist dose inducing 50% occupancy. Note that the data for intravenous (i.v.) injection (gray) refers to a previous study [17].

### Oral DCZ administration selectively induces longer-lasting behavioral effects in hM4Di-expressing monkeys

We next examined the efficacy and time-course of the chemogenetic effect of oral DCZ administration using hM4Di-expressing monkeys. We used two monkeys with hM4Di expressed in the bilateral dorsolateral prefrontal cortex (dlPFC) around the principal sulcus (Brodmann’s area 46; Fig. 3a), a brain region known to be responsible for spatial working memory and executive function [25]. Our previous studies have shown that chemogenetic silencing of the dlPFC via intramuscular DCZ administration (100 μg/kg) in these monkeys impaired performance in the spatial delayed response task (Fig. 3b), in which spatial working memory is essentially involved [17,21].

**Figure 3.**
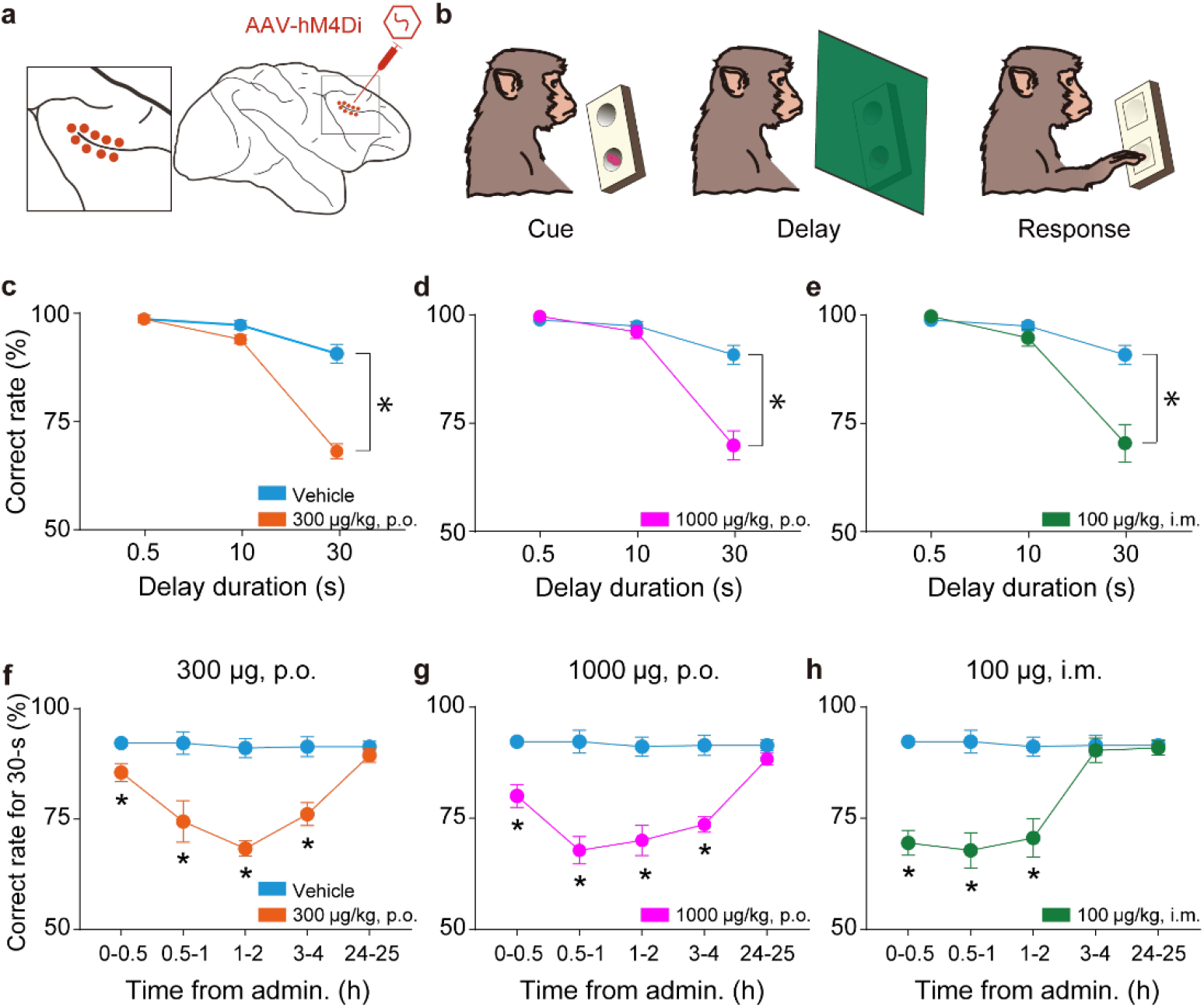
Effects of oral and intramuscular administration of DCZ on a cognitive task in monkeys expressing hM4Di in the dlPFC. **a**, Illustration showing the injection sites for AAV vector carrying an hM4Di gene. The left panel presents a zoomed-in view of the framed area of the right panel. **b**, Delayed response task. **c-e**, Effects of oral administration of 300 μg/kg (orange) and 1,000 μg/kg (magenta), and intramuscular administration of 100 μg/kg (green) of DCZ on task performance. The same data for vehicle were plotted in each panel (cyan). The behavioral tests were conducted 1-2 h following each administration. In all conditions, two-way ANOVA (treatment × delay) revealed significant main effects of treatment and delay, and interaction (100 μg/kg, i.m.; treatment: F_(1,30)_ = 17.4, p = 2.3 × 10^−4^; delay: F_(2,30)_ = 41.6, p = 2.2 × 10^−9^; interaction: F_(2,30)_ = 13.5, p = 6.4 × 10^−5^; 300 μg/kg, p.o.; treatment; F_(1,30)_ = 60.9, p = 1.1 × 10^−8^; delay; F_(2,30)_ = 117.1, p = 7.0 × 10^−14^; interaction; F_(2,30)_ = 40.5, p = 3.0 × 10^−9^; 1000 μg/kg, p.o.; treatment; F_(1,30)_ = 22.9, p = 4.2 × 10^−5^; delay; F_(2,30)_ = 62.7, p = 1.9 × 10^−11^; interaction; F_(2,30)_ = 21.4, p = 1.7 × 10^−6^), indicating the delay-dependent behavioral deficits. The data from two monkeys were pooled (N = 6 sessions; 3 sessions for each monkey). (**f-h**) Time-course of behavioral effects of each administration. Only the correct rates of 30-s trials were plotted, as they were most prominent, as shown in (**c-e**). Asterisks indicate significant differences compared to vehicle administrations (p < 0.05, Welch’s t-test, uncorrected).

We examined the behavioral effects of two oral DCZ doses (300 and 1,000 μg/kg) and an intramuscular DCZ dose (100 μg/kg) for comparison. Consistent with previous studies [17,21], intramuscular 100 μg/kg DCZ administration resulted in a delay-dependent decrease in the task performance for 1-2 h following administration (Fig. 3c). Oral DCZ administration of 300 and 1,000 μg/kg DCZ also induced a comparable impairment of the task performance for 1-2 h after administration (Figs. 3d and e). We further assessed the time-course of the chemogenetic effects by conducting a behavioral examination at 5 different time windows after treatment: 0–0.5, 0.5–1, 1–2, 3-4, and 24-25 h. We focused on the performance of the trials with the longest delay period, where the effects were most prominent (Figs. 3c-e, 30s). In the first 30-min period, intramuscular administration significantly decreased correct rates (Fig. 3f, green) compared to those after vehicle administration (Fig. 3f, cyan), indicating rapid behavioral effects as previously demonstrated [17]. Significant effects were also observed following oral administration of 300 and 1,000 μg/kg (Figs. 3f, g, orange and magenta, respectively). Similarly, all treatments had significant effects at 0.5–1 and 1–2 h after administration (Fig. 3g, h). At 3-4 h after administration, intramuscular administration had no influence on performance (Fig. 3f), while the oral administrations remained effective (Figs. 3g, h). There results were consistent between the two monkeys (Fig. S1).

To determine the minimum effective dose of DCZ via oral administration, we also administered 100 μg/kg DCZ orally and examined the monkeys 1–2 and 3–4 h thereafter (Fig. S2). The performance was significantly impaired after 1-2 h, but not after 3-4 h, indicating that this dose is effective only for a shorter time window compared to 300 and 1,000 μg/kg.

Importantly, the decrease in task performance was found to be dependent on delay duration (Fig. 3c), suggesting that the orally administered DCZ resulted in loss of dlPFC function, but did not produce side-effects on other functions, such as general attention or motivation. To further confirm that oral administration had no unwanted side effects, we conducted control experiments with the delayed response task and a simple instrumental task (Fig. S3c) in two monkeys that had not been introduced DREADDs (non-DREADD monkeys). Similar to our previous study that tested with 100 μg/kg of intramuscular DCZ administration [17], oral DCZ administration at 1,000 μg/kg dose did not produce any discernible behavioral changes in either task (Fig. S3a, d), suggesting that oral DCZ administration itself is behaviorally inert at a dose of 1,000 μg/kg or less.

Together, these results suggest that oral DCZ administration can induce behavioral effects selectively in hM4Di-expressing monkeys for hours without adverse side effects; specifically a dose of 300–1,000 µg/kg is effective for at least 4 h.

### Chronic DCZ administration enables successive induction of chemogenetic silencing for weeks

Finally, we investigated the efficacy of chronic DCZ administration across weeks. One monkey expressing hM4Di in the dlPFC was tested daily with the delayed response task to examine the chemogenetic effect of repetitive oral DCZ administration. Chronic oral DCZ administrations for at least 2 weeks was preceded and followed by several days of control vehicle administration (Fig. 4a,b), and repeated twice with a sufficient interval of several months. This design allowed us to investigate whether repetitive DCZ dose would consistently induce behavioral deficits for weeks and how soon task performance would be recovered after withdrawal of DCZ. Daily oral DCZ administrations (300 µg/kg) consistently induced an impairment of the performance of the delayed response task during 1– 2 h post-administration compared to that of following vehicle injections throughout the repetitive oral DCZ period (Fig. 4c), indicating that apparent desensitization did not occur. Previous studies have shown that chronic chemogenetic manipulation caused posttreatment effects, such as rebound excitability [3,7], suggesting that it could induce some physiological changes lasting even after agonist withdrawal. That was not the case in the current study: we found that performance recovered to a normal level on the first day after withdrawing from chronic DCZ administration (Fig. 4c), indicating that successive behavioral deficits were the outcome of temporal chemogenetic silencing rather than due to physiological change such as DREADD-induced plasticity.

**Figure 4.**
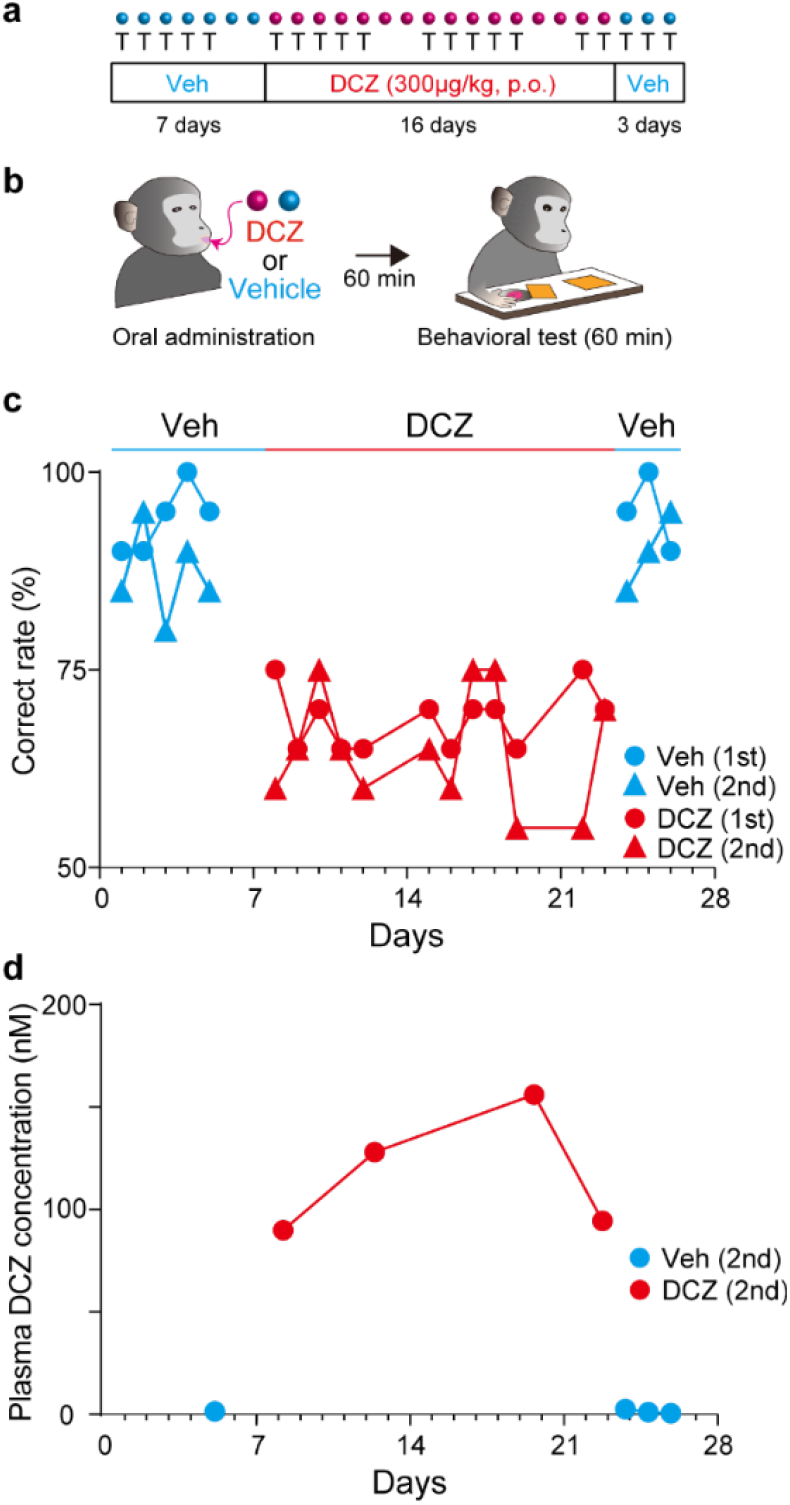
Chronic behavioral effect of daily DCZ administration across weeks. **a**, Schedule of experiments. In a series of sessions, after 7 days of vehicle administrations (Veh), DCZ was orally administered for 16 days, followed by 3 days of vehicle, once each day. The monkey was tested by the delayed response task at 5, 12 and 3 days, respectively. The series of testing were repeated twice. Cyan and purple circles represent the drugs administered (vehicle or DCZ), and the character “T” below the circles represent the days on which the behavioral tests were conducted. **b**, Schedule of an experiment in one day. **c**, Correct rates plotted as a function of days for vehicle (cyan) and DCZ (red) administrations for the 1st (circle) and 2nd (triangle) series, respectively. There was significant difference between vehicle and DCZ administrations (two-way ANOVA with treatment and schedule (1st and 2nd schedules), main effect of treatment, F_(1,36)_ = 185.6, p = 8.7 × 10^−16^). There was no significant difference between performance during the former and latter periods (two-way ANOVA with timing (former vs latter) and schedule, main effect of timing, F_(1,20)_ = 0.03, p = 0.87), indicating that the impairment was consistent during the period of administration. **d**, Plasma DCZ concentration plotted as a function of days for vehicle (cyan) and DCZ (red) administrations for monkey #245. Data were collected as with the behavioral data in (**c**), that is, 1 h after oral administration of 300 μg/kg DCZ.

It has been shown that plasma concentrations of the DCZ analogue clozapine vary within subjects over weeks of treatment [26], especially dropping by an average of more than 50% in the first 12 weeks [27]. We confirmed that the concentration of DCZ following daily administration was maintained at a similar level throughout the period of DCZ administration, indicating that chronic manipulation did not alter the pharmacokinetics of DCZ for at least 2 weeks (Fig. 4d). We also verified that chronic DCZ administration alone had no significant impact on spontaneous movements in a non-DREADD monkey (Fig. S2b). Taken together, these results suggested that DCZ is suitable for a chronic and reversible manipulation of neuronal activity, leading to potential therapeutic applications in the future.

## Discussion

Here we demonstrated that oral administration of DCZ is an effective and non-invasive means for manipulating neuronal activity through hM4Di receptors in macaque monkeys. Our pharmacokinetics and *in vivo* PET occupancy examination showed that the bioavailability of orally administered DCZ was about 10-20% of that of intramuscular injection, and thus the appropriate dosage of oral DCZ to exert hM4Di-mediated chemogenetic effects was estimated to be 300–1,000 μg/kg. Indeed, this oral DCZ at this range of doses caused severe impairments in working memory performance for several hours in monkeys with hM4Di expressed in dlPFC, but did not induce any discernible behavioral side-effects in control non-DREADD-expressing monkeys. Furthermore, repetitive daily oral DCZ doses over two weeks yielded consistent chemogenetic effects without apparent sign of desensitization. Taken together, the oral administration of DCZ affords a minimally invasive strategy for chronic and reversible chemogenetic control of neuronal activity via muscarinic DREADDs in nonhuman primates, thereby providing a great potential for its clinical application.

For DREADDs studies in rodents and nonhuman primates, systemic (intraperitoneal, intramuscular, or intravenous) injections are the standard approach of agonist administration, as they allow precise control of dosage and timing. However, such approach forces the animals to be restrained, which may cause stress responses and unwanted effects on behavioral actions. Several DREADDs studies have demonstrated that oral administration seems effective for overcoming this issue. For example, oral CNO administration induced changes in ethanol consumption by inhibition of the nucleus accumbens via hM4Di, in which neuronal activity was remotely manipulated while the animals were unrestrained [28]. This is beneficial when wanting to examine the effect on natural behavior including social communication [29]. Oral CNO administration has also been applied to manipulate the activity of non-neuronal cells such as microglia [5]. However, recent reports have revealed that the effects of CNO are mediated by its back-conversion to clozapine which crosses the blood-brain barrier and acts as a DREADDs actuator [13]. Oral administration of low doses of clozapine and olanzapine as agonists has also been attempted [7]. Due to their high affinities for endogenous receptors, however, possible side effects are always a matter of concern that cannot completely be ruled out.

In the current study, we evaded these issues by using DCZ, a novel DREADDs agonist with high brain permeability and high selectivity. Since the effects of drugs on the central nervous system depend on multiple factors such as brain permeability, drug kinetics, and affinity for target receptors, it is generally difficult to predict the optimal drug dosage. As we have demonstrated in previous studies [17,19], PET occupancy measurements together with pharmacokinetics analysis are extremely effective ways for seeking the adequate agonist dose range. We found that the bioavailability of orally administered DCZ was 10-20% compared to systemic injections (see Figs. 1 and 2). Based on this estimation, the dose range suitable for hM4Di activation was determined to be 300-1,000 μg/kg. In line with the kinetics data, such oral DCZ doses induced relatively long-lasting effects compared to intramuscular injections. The DCZ kinetics and behavioral effects were examined up to 4 h after oral administration, during which the concentration of DCZ persisted in CSF, and thus the chemogenetic effects may be maintained longer. Considering potential side-effects due to action on endogenous receptors as indicated by other prior DREADDs agonists, we performed control experiments using non-DREADD expressing monkeys, demonstrating that these oral DCZ doses alone did not produce any discernible changes in behavior associated with spatial working memory or reward expectation (Fig. S2).

In the present study, we did not examine the effects of oral DCZ administration on hM3Dq, an excitatory muscarinic DREADDs that responds to DCZ [17]. Our previous work showed that the lower doses (1-3 μg/kg) of DCZ through intramuscular injection were capable of inducing significant neuronal excitation in mice and monkeys expressing hM3Dq [17]. Thus, 3-30 μg/kg would be an effective range for activating hM3Dq via oral administration. Indeed, we have recently demonstrated that marmosets expressing hM3Dq produced consistent behavioral changes when fed a diet containing 10 μg/kg of DCZ [18]. In any case, it should be noted that the optimal agonist dose generally depends on the level of the overexpressed functional protein and the targeted circuit to be manipulated [30]. In addition, as is usual in studies using genetic methods, control experiments (e.g., using non-DREADDs animals) are recommended.

It has been shown that chronic administration of DREADDs agonists induces DREADDs-mediated changes for days to weeks in rodents [3,4,6,7]. However, some studies reported that the effects following administrations of agonists were not consistent during treatments. For example, Goossens et al. reported that the administrations of clozapine and olanzapine caused significant behavioral changes for several days, but subsequently such effects diminished even though the agonist administrations continued [7]. In addition, it has been reported that inhibition by hM4Di with agonist treatment over weeks induced posttreatment rebound excitability [3,7]. In contrast to these studies, daily oral DCZ delivery constantly impaired the monkey’s performance throughout the administration periods (see Fig. 4). Moreover, performance on the day following completion of repeated doses of DCZ was as high as the baseline, suggesting that the weeks-long inactivation of the PFC did not affect subsequent behavior. These results suggest that chronic DCZ administration did not cause desensitization of hM4Di or any long-lasting change, such as plasticity of neural circuits that govern working memory. It remains to be clarified whether this is a general capability of DREADD/DCZ or is due to the characteristics of neurons and local circuits in the dlPFC as a target. Irrespective of the mechanism, the present results indicate that it is possible to investigate the effects of chronic attenuation of PFC activity on various cognitive functions. Importantly, it can resolve potential discrepancies that may arise in the behavioral outcomes of different durations or methods of inactivation. For example, it has been demonstrated that acute and chronic silencing of PFC interneurons had opposite impacts on anxiety-like behavior in mice [31], suggesting that short-vs. long-term manipulations of local circuits may differentially alter PFC network functions. Besides, long-term abnormalities or imbalances in prefrontal activity have been suggested as a pathophysiological mechanism for psychiatric disorders, since chronic stress exposure could lead to anxiety disorder and depression [32]. DREADDs subserve to mimic or reverse such a long-term circuitry, changes that are beyond the reach of conventional acute blockade and/or irreversible lesioning. The chemogenetic approach introduced in the present study will expand the opportunity of acute and chronic manipulations of specific circuits in the primate prefrontal cortex, thereby leading to a better understanding of higher brain functions in health and disease conditions. In addition, it has the potential to be used in therapeutic applications for symptoms caused by abnormal neuronal activity in a specific brain region, such as seizures in epilepsy.

Finally, it should be noted that this study did not examine continuous chemogenetic effects beyond the duration of action of a single oral dose (∼4 h). Continuous silencing may be achieved with more frequent and repeated administrations. Future studies will need to determine the dosage and interval of administrations to maintain sufficient DCZ concentration to attain continuous chemogenetic effects, and to ensure that it does not cause undesirable side effects, toxicity or tachyphylaxis.

DREADDs technology is becoming increasingly popular as a means to control neuronal activity remotely, less-invasively and reproducibly in rodents and nonhuman primates. With the accumulation of numerous successful reports, muscarinic DREADDs are now under consideration for clinical application. Given the long-term stable chemogenetic effects with orally delivered DCZ in macaque monkeys, it holds great promise for the translational use of DREADDs technology, especially in the development of therapeutic trials for neurological and neuropsychiatric disorders.

## Materials and Methods

### Subjects

A total of 8 macaque monkeys [7 Japanese (*Macaca fuscata*) and 1 Rhesus monkeys (*Macaca mulatta*); 6 males, 2 females; 2.8-8.0 kg; age 4-9 years at the beginning of experiments] were used (Table S1). The monkeys were kept in individual primate cages in an air-conditioned room. A standard diet, supplementary fruits/vegetables and a tablet of vitamin C (200 mg) were provided daily. All experimental procedures involving animals were carried out in accordance with the Guide for the Care and Use of Nonhuman primates in Neuroscience Research (The Japan Neuroscience Society; https://www.jnss.org/en/animal_primates) and were approved by the Animal Ethics Committee of the National Institutes for Quantum and Radiological Science and Technology.

### Viral vector production

Adeno-associated virus (AAV) vector expressing hM4Di (AAV1-hSyn-hM4Di-IRES-AcGFP, 4.7 × 10^13^ particles/mL) was produced by helper-free triple transfection procedure, which was purified by affinity chromatography (GE Healthcare, Chicago, USA). Viral titer was determined by quantitative PCR using Taq-Man technology (Life Technologies, Waltham, USA).

### Surgical procedures and viral vector injections

Surgeries were performed under aseptic conditions in a fully equipped operating suite. We monitored body temperature, heart rate, SpO_2_ and tidal CO_2_ throughout all surgical procedures. Monkeys were immobilized by intramuscular (i.m.) injection of ketamine (5-10 mg/kg) and xylazine (0.2-0.5 mg/kg) and intubated with an endotracheal tube. Anesthesia was maintained with isoflurane (1-3%, to effect). Before surgery, magnetic resonance (MR) imaging (7 tesla 400mm/SS system, NIRS/KOBELCO/Brucker) and X-ray computed tomography (CT) scans (Accuitomo170, J. MORITA CO., Kyoto, Japan) were performed under anesthesia (continuous intravenous infusion of propofol 0.2-0.6 mg/kg/min). Overlay MR and CT images were created using PMOD® image analysis software (PMOD Technologies Ltd, Zurich, Switzerland) to estimate stereotaxic coordinates of target brain structures.

One monkey (#237) received injections of AAV vector into one side of the amygdala (2 µL x 2 locations). The monkey underwent a surgical procedure to open burr holes (∼8-mm diameter) for the injection needle. Viruses were pressure-injected using a 10-μl Hamilton syringe (model 1701 RN, Hamilton) with a 30-gauge injection needle and a fused silica capillary [19]. The Hamilton syringe was mounted into a motorized microinjector (UMP3T-2, WPI) that was held by a manipulator (model 1460, David Kopf) on the stereotaxic frame. After the dura mater was opened about 3mm, the injection needle was inserted into the brain and slowly moved down 2 mm beyond the target and then kept stationary for 5 min, after which it was pulled up to the target location. The injection speed was set at 0.25 μl/min. After each injection, the needle remained in situ for 15 min to minimize backflow along the needle.

Two monkeys (#229 and #245) received injections of AAV vector into the bilateral prefrontal cortex (Brodmann’s area 46) as described previously [17]. The frontal cortex was exposed by removing a bone flap and reflecting the dura mater. Handheld injections at an oblique angle to the brain surface were made under visual guidance through an operating microscope (Leica M220, Leica Microsystems GmbH, Wetzlar, Germany). Nine tracks were injected in each hemisphere located at the caudal tip (1 track) and along the dorsal (4 tracks) and ventral (4 tracks) bank of the principal sulcus. Viral vectors were injected at 3 to 5 µL per track depending on the depth. A total of 35 to 44 µL of viral aliquots was injected into each hemisphere.

### Drug administration

DCZ (MedChemExpress HY-42110) was dissolved in 2.5% dimethyl sulfoxide (DMSO, FUJIFILM Wako Pure Chemical Co.), aliquoted and stored at –30°. For systemic i.m. injection, this stock solution was diluted in saline to a final volume of 100 μg/kg, and injected intramuscularly. For systemic oral administration (per os, p.o.), the DCZ stock solution was diluted in saline to a final volume of 100 μg/kg, 300 μg/kg, or 1,000 μg/kg. The solution was then injected into some pieces of snack or diluted in drinking water, which were given to the monkeys. It typically took 2-3 min for the monkeys to consume. After each administration, we confirmed that there was no food in the monkeys’ mouths. Fresh solution was prepared on each day of usage.

### Occupancy study

An occupancy study was performed using one monkey expressing hM4Di in the amygdala (#237). PET scans were performed using a microPET Focus 220 scanner (Siemens Medical Solutions USA, Malvern, USA). Monkeys were immobilized by ketamine (5-10 mg/kg) and xylazine (0.2-0.5 mg/kg) and then maintained under anesthetized condition with isoflurane (1-3%) during all PET procedures. Transmission scans were performed for about 20 min with a Ge-68 source. Emission scans were acquired in 3D list mode with an energy window of 350–750 keV after intravenous bolus injection of [^11^C]DCZ (324.9-382.3 MBq). Emission data acquisition lasted for 90 min. Pretreatment of DCZ or vehicle p.o. via a gastric catheter was followed by anesthesia and emission scans at 30 and 90 min, respectively. To estimate the specific binding of [^11^C]DCZ, regional binding potential relative to nondisplaceable radioligand (BP_ND_) was calculated by PMOD® with an original multilinear reference tissue model (MRTMo)[20]. The volume of interest for the hM4Di-positive region was defined as the area where the BP_ND_ value of [^11^C]DCZ was higher than 0.5, while that of the control region was placed at the corresponding contralateral side.

Estimates of fractional occupancy were calculated according to the previous study [17] by the following equation:

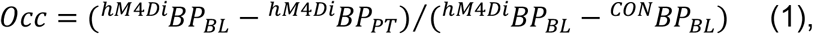

where ^*hM4Di*^*BP*_*BL*_ and ^*CON*^*BP*_*BL*_ indicate BP_ND_ at the hM_4_Di-expressing amygdala region and contralateral control region under baseline conditions, respectively, while ^*hM4Di*^*BP*_*PT*_ indicates BP_ND_ at the hM_4_Di-expressing amygdala region under pretreatment condition. The relationship between occupancy (*Occ*) and agonist dose (*D*_*DCZ*_) was modeled by the Hill equation

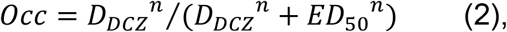

where *ED*_*50*_ and *n* indicate the agonist dose achieving 50% occupancy and the Hill coefficient, respectively.

### Pharmacokinetics analysis

Three monkeys were used to assess the concentration of DCZ and its metabolites in plasma or cerebrospinal fluid (CSF). Blood and CSF were collected at 30, 60, 90, 120, 150, 180, 210, 240 min, and 24 h after DCZ administration (100 μg/kg, i.m. or 300 μg/kg, p.o.) under isoflurane (1-3%, to effect) anesthesia. Blood was collected with a heparinized syringe and plasma samples was obtained by centrifugation at 3,500 g for 15 min. All samples were stocked at -80°C until analysis.

The pretreatment protocols for CSF and plasma samples were described previously [17]. Quantification of DCZ, and its metabolites C21 and DCZ-N-oxide, was performed by multiple reaction monitoring (MRM) using a Shimadzu UHPLC LC-30AD system (Shimadzu Corp., Kyoto, Japan) coupled to a tandem MS AB Sciex Qtrap 6500 system (AB Sciex LLC, Framingham, USA). The following MRM transitions (Q1/Q3) were used to monitor each compound: DCZ (293.0/236.0), C21 (279.0/193.0), DCZ-N-oxide (309.0/192.9). Other protocols were described previously.

### Behavioral task

Three monkeys (#226, #229, and #245) were tested with a spatial delayed response task. The protocol was almost the same as of our previous study [21]. Behavioral testing was conducted in a sound-attenuated room. Monkeys sat on a monkey chair from which they could reach out one hand and take food to their mouths. A wooden table with two food-wells was placed in front of the monkeys, and a screen was placed between the monkeys and the table. First, a piece of food reward (raisin or snack) was placed in one of the two food-wells, and then both wells were covered with wooden plates. Then, the screen was placed for 0.5, 10 or 30 s, which served as delay periods. The position of the baited well (left or right) and the delay period (0.5, 10 or 30 s) were determined pseudo-randomly. After the delay period, the screen was removed and the monkeys were allowed to select either food-well. The monkeys were allowed to get the food if they reached for the correct food-well and removed the cover plate. The inter-trial interval was set at 10 s. A daily session lasted about one hour, and consisted of 3 blocks of 30 trials for monkey #229, and 2 blocks of 30 trials for monkeys #226 and #245, which were interleaved with 5-min rest periods. Behavioral tests began immediately or 0.5, 1, 3, or 24 h after vehicle or DCZ treatment.

Another monkey without AAV injection (non-DREADD; #239) was tested with a reward-size task using the same protocol as applied in a previous study [22]. The behavioral testing began 1 h after an oral administration of either vehicle or DCZ.

### Statistics

To examine the effect of each treatment on the performance of the delayed response task, behavioral measurement (correct rates) was subjected to Welch’s t-test or two-way ANOVA using GraphPad Prism 7.

## Acknowledgments

We thank J. Kamei, R. Yamaguchi, Y. Matsuda, Y. Sugii, T. Okauchi, T Kokufuta, M. Fujiwara and M. Nakano for their technical assistance. Monkeys used in this study were partly provided by National Bio-Resource Project ‘‘Japanese Monkeys’’ of MEXT, Japan.

---

**Table S1.** Summary of monkeys used in the study.

| ID | Species | Sex | Age (y) | Vector | Occupancy | PK | Behavior |
| --- | --- | --- | --- | --- | --- | --- | --- |
| 226 | J | F | 8 | N/A |  | ✓ | ✓ |
| 229 | R | M | 5 | hM4Di<br>dIPFC |  |  | ✓ |
| 237 | J | M | 7 | hM4Di<br>amygdala | ✓ |  |  |
| 239 | J | M | 6 | N/A |  |  | ✓ |
| 245 | J | F | 7 | hM4Di<br>dIPFC |  | ✓ | ✓ |
| 248 | J | M | 4 | N/A |  | ✓ |  |
| 249 | J | M | 4 | N/A |  | ✓ |  |
| 253 | J | M | 5 | N/A |  |  | ✓ |
| Total | 8 (R,1/J,7) | F,2; M,6 |  | 3 | 1 | 4 | 5 |

A tick indicates the subject was used in the experiment. PK, pharmacokinetics study; Species (J, Japanese; R, Rhesus), Sex (F, Female; M, Male), Age (years).

**Figure S1.**
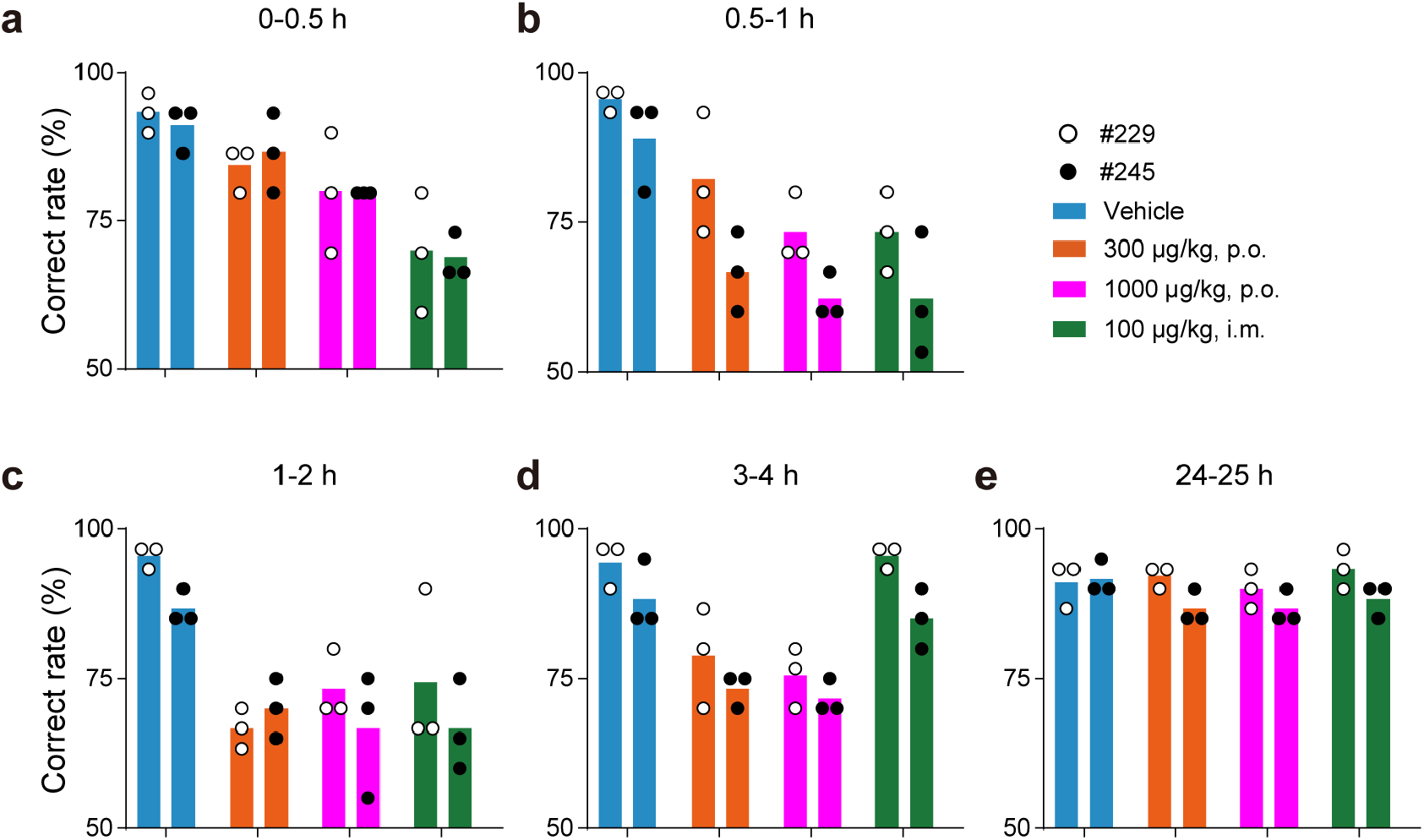
Consistent behavioral effects of DCZ administration between two monkeys expressing hM4Di in the dlPFC. **a-e**, Correct rate of the longest delay (30s) trials in delayed response task following oral administration of vehicle (cyan), 300 μg/kg (orange) and 1000 μg/kg (magenta), and intramuscular administration of 100 μg/kg DCZ (green) for each monkey (open and filled circles for #229 and #245, respectively) in different time windows: 0-0.5 h (**a**), 0.5-1 h (**b**), 1-2 h (**c**), 3-4 h (**d**), and 24-25 h post administration (**e**), respectively. Note that Fig. 3 shows the same data pooled together.

**Figure S2.**
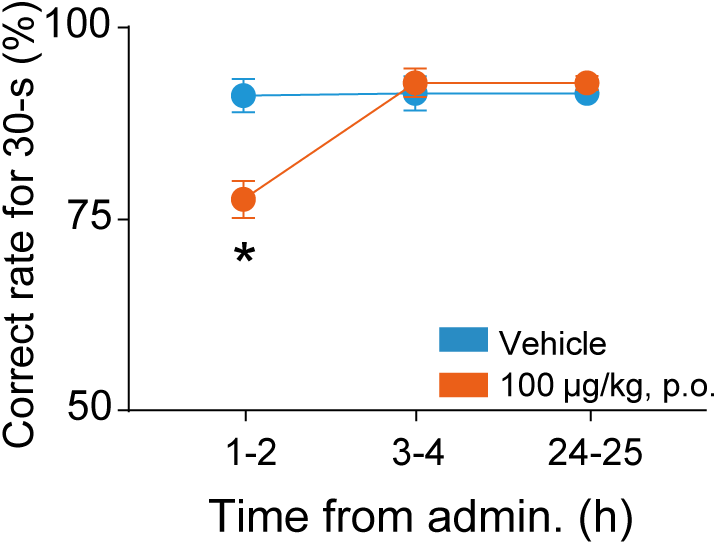
The time course of the chemogenetic effects of oral administration of 100 μg/kg DCZ on the delayed response task. The behavioral tests were conducted 1-2, 3-4, and 24-25 h following each administration. There was significant difference between vehicle and 100 μg/kg oral DCZ administration at 1-2 h, but not at 3-4 and 24-25 h (p < 0.05, Welch’s t-test, uncorrected). Only the data for 30-s delay trials were plotted.

**Figure S3.**
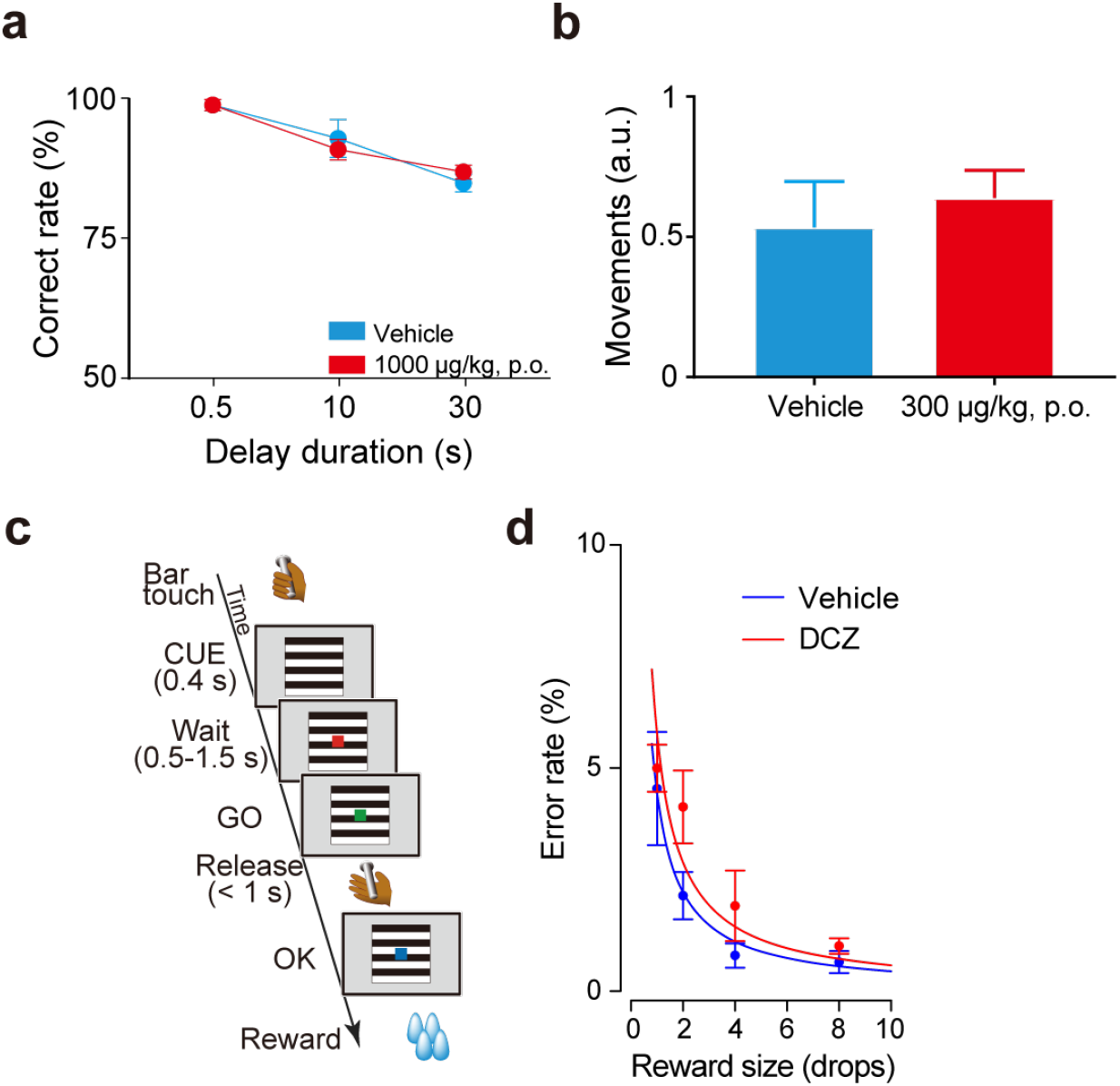
Effects of 1000 µg/kg DCZ oral administration on behavioral tasks in non-DREADD animals. **a**, Performance of delayed response task following oral administration of vehicle (cyan) and 1000 μg/kg DCZ (red) in a non-DREADD animal (#226). There was no significant difference between treatments (two-way ANOVA, treatment × delay, treatment: F(1,24) = 0, p = 1). **b**. Effects of chronic DCZ administration on spontaneous movement in a DREADD-expressing animal (#253). **c**, Illustration of a reward-size task. Each trial in this task began when the monkey touched a lever, which was followed by the appearance of a visual cue signaling the size of the upcoming reward (1, 2, 4 or 8 drops). To obtain the reward, the monkeys had to release the lever when a visual target changed color from red to green. **d**, Example behavioral data of reward size task for DCZ (1000 µg/kg, p.o.; red) and vehicle administration session (cyan) obtained from a non-DREADD monkey (#239). Two-way ANOVA (treatment × reward size) revealed no significant difference (p > 0.05).

